# Targeting Aryl Hydrocarbon Receptor with small molecule, 1′H-indole-3′-carbonyl-thiazole-4-carboxylic acid methyl ester blocked human glioma cell invasion via MYH9

**DOI:** 10.1101/2020.01.13.903674

**Authors:** Lijiao Zhao, Qiuting Shu, Hui Sun, Yunlong Ma, Dandan Kang, Yating Zhao, Jing Lu, Pei Gong, Fan Yang, Fang Wan

**Affiliations:** Inner Mongolia Agricultural University

**Keywords:** AHR, ITE, Glioma, MYH9, Mesenchymal migration, Amoeboid migration

## Abstract

Aryl hydrocarbon receptor (AHR) was a master regulator of anti-tumor cell migration in various cell types. Whether and how AHR regulates glioma cell migration is largely unknown. We found that small molecule 2-(1′H-indole-3′-carbonyl)-thiazole-4-carboxylic acid methyl ester (ITE), an endogenous AHR ligand, can significantly block glioma cell migration and invasion *in vitro*, *ex vivo* and *in vivo*. Knocking down AHR by siRNA abolished ITE’s migration-inhibiting effects. ITE increased the number of filopodia-like protrusion formation, but reduced protrusion attachment to the extracellular matrix, and inhibited the rear retraction of migrating glioma cells. Moreover, both mesenchymal and amoeboid migrating cells were observed in the DMSO control group while none of the cells display amoeboid migration in the ITE treated group. MYH9 was significantly reduced by ITE treatment in human glioma cells. Over-expression of MYH9 abrogated ITE’s migration-inhibiting effects, with the expression level of MYH9 correlated with cell migration ability. Since MYH9 is a component of non-muscle myosin IIA (NMIIA), which is essential for cell migration in 3D confined space, and not a discovered target of AHR, the fact that ITE affects MYH9 via AHR opens a new research and development avenue.

## Introduction

Glioblastoma (GBM), the most common and advanced form of glioma, is one of the most lethal human cancers. GBM patients survive, on average, 12–15 months despite aggressive surgical resection, radiation, and chemotherapy[1]. The diffuse invasion of the GBM tumor cells and the blood-brain barrier pose major challenges for successful therapy since invaded tumor cells are sheltered from complete surgical removal and radiation. Targeted drug discovery concentrates on the receptor and signaling pathway, such as EGFR, VEGFR, PDGFR, MET, MEK, and molecules that mediate cell interaction with the ECM, such as integrin and syndecan.

We targeted AHR, a ligand-activated transcription factor, and E3 ubiquitin ligase. AHR regulates cytotoxic immune cells in cancer by controlling the function and differentiation of multiple types of immune cells. In glioma, AHR is expressed in glioma and the stromal cells, including endothelial cells, astrocytes, oligodendrocytes, microglia, macrophages, dendritic cells and T lymphocyte[2–4]. AHR’s effects on cancer are complex, highly depends on the ligands in the microenvironment and the cell types. Accumulating evidence support AHR’s role as a tumor suppressor in glioma[5].

We found that targeting AHR in glioma blocked glioma invasion. The majority of endogenous AHR ligands are tryptophan metabolites, and the most abundant tryptophan metabolite in glioma is kynurenine, which is produced by enzymes 2,3-dioxygenase (IDO/TDO), binds to AHR and inhibits cytotoxic immune cells activity against glioma cells[6], forming a pathway of Tryptophan-Kynurenine-AHR-Impairment of Immunity. To block this pathway, inhibitors for the IDO/TDO were developed and mixed clinical trial results have been posted[7, 8], not meeting expectations. An alternative route is to directly target AHR using small-molecule ligands. We hypothesized that targeting AHR using a high-affinity agonist could achieve this goal while maintaining AHR’s tumor suppressor effects in glioma. To this end, we tested the effects of ITE, an endogenous AHR agonist, and discovered that ITE significantly inhibited glioma cell migration and invasion.

We report that ITE can significantly block glioma cell migration and invasion in wound healing and transwell assay (*in vitro*), in brain invasion assay (*ex vivo*), and in a mouse model (*in vivo*). In a 3D collagen matrix, in addition to the more dynamic filopodia, ITE induced weaker protrusion attachment to the ECM. In 2D cultures, glioma cells failed to undergo amoeboid migration when treated with ITE and subject to pH changes. ITE blocked the invasion of tumor into the surrounding brain parenchyma, but not that toward the ventricles. ITE has a higher efficacy binding to AHR than previously reported ligands and possibly using a different mechanism for action. MYH9, a component of NMIIA, is altered in ITE treated mouse and essential for ITE action.

## Materials and Methods

### Cell culture

Human and mouse glioma cell lines U87MG and GL261 were obtained from the National Infrastructure of Cell Line Resource and the Third Military Medical University, respectively. U87MG and GL261 cells were cultured using MEM/EBSS and DMEM/F12 supplemented with 10%FBS respectively.

### Cell invasion and migration assays

Transwell chamber (CORNING, USA) was coated with 30 µl rat tail tendon collagen type I (PERFEMIKER, #C20-200110), incubated for 30 min at 37°C. 5×10^4^ cells were seeded into chambers and cultured in DMEM/F12 with 2% FBS, and the lower chamber medium contains 10% FBS. After 24hr, cells were fixed with 4% paraformaldehyde, and the non-migrated cells were removed with cotton swabs. The migrated cells on the bottom were stained with crystal violet and counted. Invasion assays were repeated six times over multiple days.

For the wound-healing assay, cells were grown to 90% confluence in 12 well tissue culture dish and scratched using a 10ul pipette tip. Cultures were rinsed with PBS and incubated in serum-free medium for 20hr. Images were captured at 0hr and 20hr, and the area of the scratch was measured using ImageJ. Two or three fields per well in three wells were quantified for each condition. DMSO(SIGMA,#D5879) was used as control.

### Mouse orthotopic glioma model

All experiments were performed following the animal care guidelines of Inner Mongolia Agricultural University.

8-week-old C57BL/6 Mice (Beijing Vital River Laboratory Animal Technology Co. Ltd) were anesthetized with an intraperitoneal injection of 0.3% Pentobarbital sodium(30mg/kg). For the stereotactic intracranial injection, the surgical site was shaved and prepared with iodine. A midline incision was made to expose the bregma point, and a 1 mm diameter right parietal burr hole was drilled with a dental drill, centered at 2 mm posterior to the coronal suture and 2 mm lateral to the sagittal suture. Mice were placed in a stereotactic frame and 1×10^5^ GL261 cells in 2.5µL medium were intracranially-injected at a depth of 3 mm using Microinjection Pump (KD scientific, 78-1311Q) at a speed of 0.5ul/min. The needle was removed slowly and the skin was sutured with nylon thread.

### Hematoxylin and eosin stain of the mouse brain tissue

Mice were anesthetized and perfused with 4% paraformaldehyde (PFA) through the right atrium. Brains were harvested and placed in 4% PFA for 12hr, then in 15%, 20%, 30% sucrose solutions consequently for 12hr each. Tissue was embedded in Optimal Tissue Cutting Compound (OCT# SAKURA) and frozen in liquid nitrogen. Tissue was sectioned at 12um using frozen slicer (LEICA #CM1850 Germany). Sections were then stained with hematoxylin and eosin (Sigma Aldrich, #SLBS9620), and imaged using microscope and stereomicroscope (Nikon, Tokyo, Japan).

### Organotypic Brain Slices Culture

Mouse brains were extracted from 6-week old female C57BL/6 mice (euthanized by cervical dislocation), placed in PBS solution with 3% penicillin and streptomycin in a sterile Petri dish. Three percent low melting point agarose (Solarbio, #A8350) solution was prepared and cooled to touch, then poured over the brain, and cooled for an additional 30-45 min. The agarose gel was trimmed and 300um organotypic brain slices were generated using a vibrating microtome ZQP-86 (Zhi Xin Biotech, Shang Hai, China). The brain slices were sectioned into two symmetrical halves and transferred to 6-well transwell plate chamber by spatula and placed next to each other. Two mL DMEM /F12 medium with 25% FBS was added to the lower chamber and placed in a 37°C, 5% CO_2_ incubator for overnight culture.

### Organotypic Brain Slice Invasion Assay

10^5^ GL261 cells were incubated with CM-Dil (YEASEN, # 40718ES60) for 15min in 2 µL media, seeded onto the seam where two brain slices met, and the brain slices were cultured at the air-liquid interface, with the culture media refreshed every 2-3 days. For drug treatments, serum-free media were added to the lower chamber, with different doses of ITE(0.1nM,10nM,1000nM), 3000nM PTK2 inhibitor PF431396 (MCE, # 717906-29-1) or 0.02% DMSO control. After 5 days and 8 days of continuous culture, cell invasion was observed under a stereomicroscope (Nikon, Tokyo, Japan) for overall slice imaging. For high-resolution imaging, multiple overlapping pictures were taken and stitched together using the software panoramamaker6.

### 3D culture of GL261 cells

2×10^5^ cells at 50ul were mixed with 12ul NaOH, 710ul complete medium and 23 ul PBS, and 220ul rat tail tendon collagen type Ⅰ (PERFEMIKER, #C20-200110). After solidification, supplement 1ml complete medium to the gel. Time-lapse photography was performed using an OlympusIx71 microscope with software Cell Sens Standard, at 30s intervals for 1 hour. Time-lapse videos were made using ImageJ.

### AHR siRNA gene knockdown and total RNA extraction

8×10^5^ U87MG cells were seeded into a 6-well plate and transfected when cells grow to 60% confluence according to the manufacturer’s instructions. After 4hr of transfection, fresh complete medium was added, and total RNA was extracted from the cells after 36hr transfection using RNA Fastagen Kit (Shanghai Feijie Bio, #220010) according to the manufacturer’s protocol.

### Western Blot

Protein was extracted on ice in RIPA solution (Solarbio, #R0020), and quantified using a bicinchoninic acid protein assay kit (Solarbio, #PC0020). Forty μg protein was separated by 8% SDS/PAGE, transferred to nitrocellulose membranes (MILLIPORE, #HATF00010) at 120v for 90min. Membranes were blocked with 5% Skim Milk solution (Solarbio, #D8340) for 3h at 37°C, immunoblotted with primary antibody (listed in Supplementary Table 1) overnight at 4°C, washed, then with a secondary antibody, washed, and imaged using ODYSSEY CLX (Clx-0519, LI-COR, Gene Company Limited, USA)

### Transient transfection of cells

8×10^5^ U87MG cells were plated into 6-well plates 24hr before transfection, and the cells were washed and supplemented with 2ml DMEM before transfection. Plasmid DNA and PEI (ALDRICH, Lot #BCBT0649) were each diluted with MEM/EBSS and allowed to stand for 10 min. The DNA solution was added into the PEI solution and mixed well at N/P=20, incubated for 10 min. PEI/DNA complexes were added to the cells and replaced by fresh medium after 3.5hr. The empty vector was employed as a control. MYH9 transgene expression was verified using real time-PCR.

### Real time-PCR

Four hundred ng total RNA was employed for cDNA synthesis using ProtoScript II First Strand cDNA Synthesis kit(TaKaRa,#RR036A) according to the manufacturer’s protocol.Forty ng cDNA was used as templates for RT-PCR using SYBR® Premix Ex Taq kit (TaKaRa,#RR820A) using LightCyclerW 480 (Roche, Switzerland). The sequences and efficiency of all primers were listed in (Supplementary Table 2), and HPRT/GAPDH was used as reference genes.

### Statistical analysis

All data were presented as the mean ± standard deviation, and Student’s t-test was applied assuming equal variance between groups.

## Results

### ITE inhibited glioma migration and invasion *in vitro, ex vivo* and *in vivo*

We investigated ITE’s effects on glioma cells using wound healing migration and transwell invasion assays. ITE significantly inhibited cell migration in a non-monotonic dose-response fashion (Fig. 1A-B), probably due to the response to a mixture of both ITE and other AHR ligands that exist in the culture environment. ITE significantly blocked the invasion of both U87MG and GL261 cells (Fig. 1C-D). We knocked-down AHR level in the U87MG cells by siRNA (Fig. 1E) and found that the knocking down of AHR abolished most of the ITE’s migration-inhibiting effects, indicating such effects depended on AHR (Fig. 1F).

**Fig. 1.**
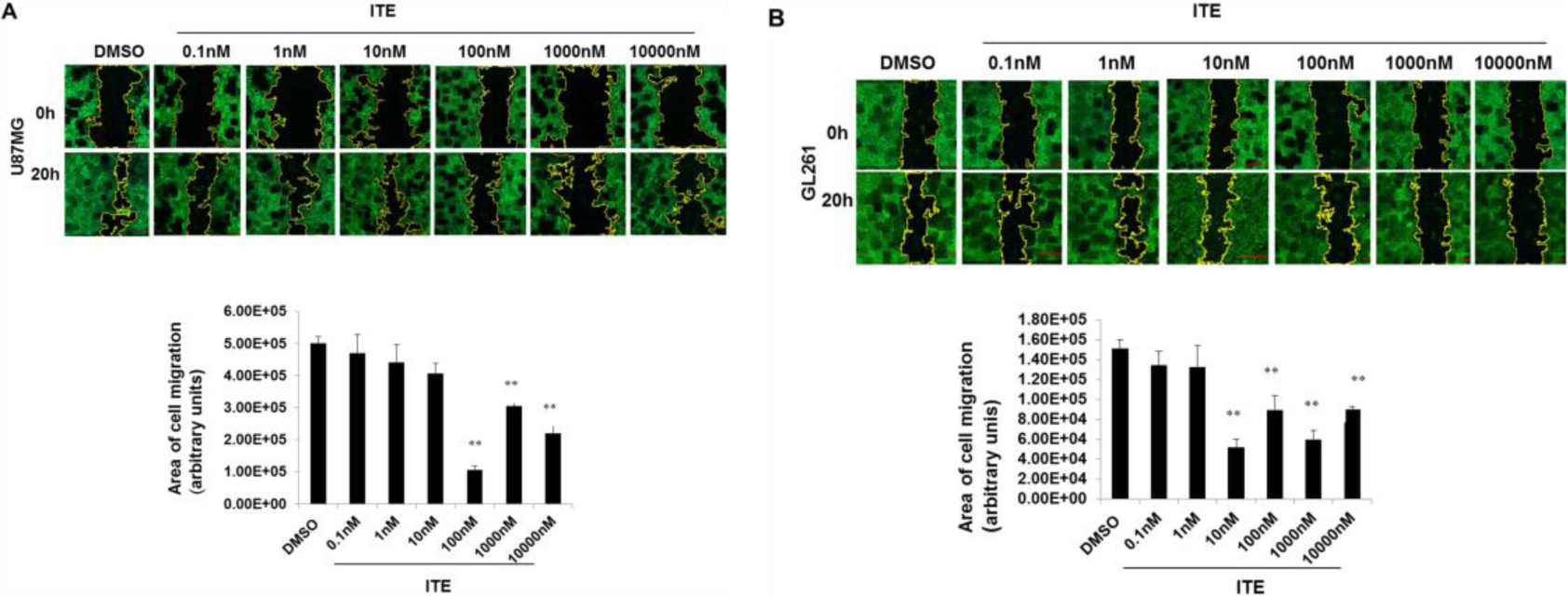
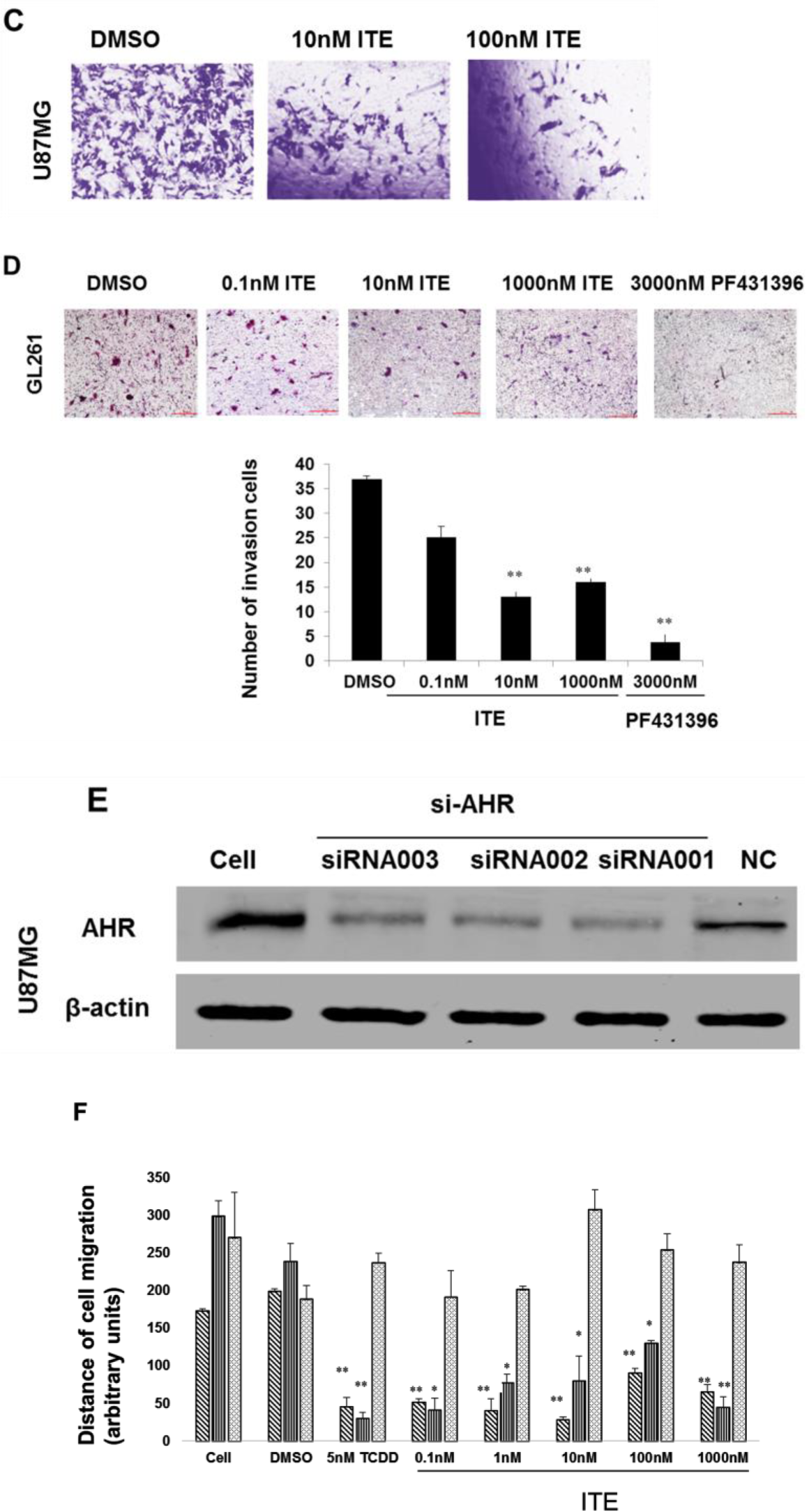
ITE inhibited migration and invasion of glioma cells in wound healing and transwell assay. (A-B) Wound healing assays. U87 and GL261 cells were treated with DSMO or various doses of ITE for 20hr. The area of the wound was measured in 3 replicate wells per treatment condition (*p<0.05**P <0.01). (C-D)Transwell assays. The cells were treated with various concentrations of ITE, DMSO or PF431396 (PTK2 inhibitor) for 20hr. Those migrated through the rat tail tendon collagen type I gel and the membrane was stained and counted. (*p<0.05 **P <0.01). (E) siRNA knocking down of AHR expression in U87MG cells. Three different AHR siRNAs or NC were transfected into the U87 cells for 36h and the AHR level was assessed by western blot. (F) AHR knockdown abolished ITE’s migration inhibition effect. U87MG cells were untransfected (AHR), transfected with AHR siRNA002, or with NC for 36hr, and treated with DMSO, various concentrations of ITE, or 5nM TCDD, for additional 18hr. Cell migration was assessed by the migration distance of three replicate wells in the wound healing assay (*p<0.05, **p<0.01).

To test whether ITE can block invasion into the brain parenchyma, we performed *ex vivo* brain invasion assay using live-cell dye CM-Dil labeled GL261 cells (Fig. 2A-B). We found that ITE significantly inhibited GL261’s invasion into the normal mouse brain parenchyma with high efficacy, showing significant inhibition at 10nM compared to PTK2 inhibitor PF431396. We tested whether ITE can block invasion in an orthotopic mouse model of glioma using GL261 cells injected into the front lobe of the mouse brain (Fig. 2C). Typical glioma invasion patterns were observed in this model, including into the peritumor parenchyma and toward the ventricles (Fig. 2D). ITE (100mg/kg body weight, given every other day) significantly blocked peritumor invasion, as the tumors of the ITE group had smoother edges compared to those of the control group (Fig. 2E). The invasions into the ventricles were not significantly altered after ITE treatment.

**Fig. 2.**
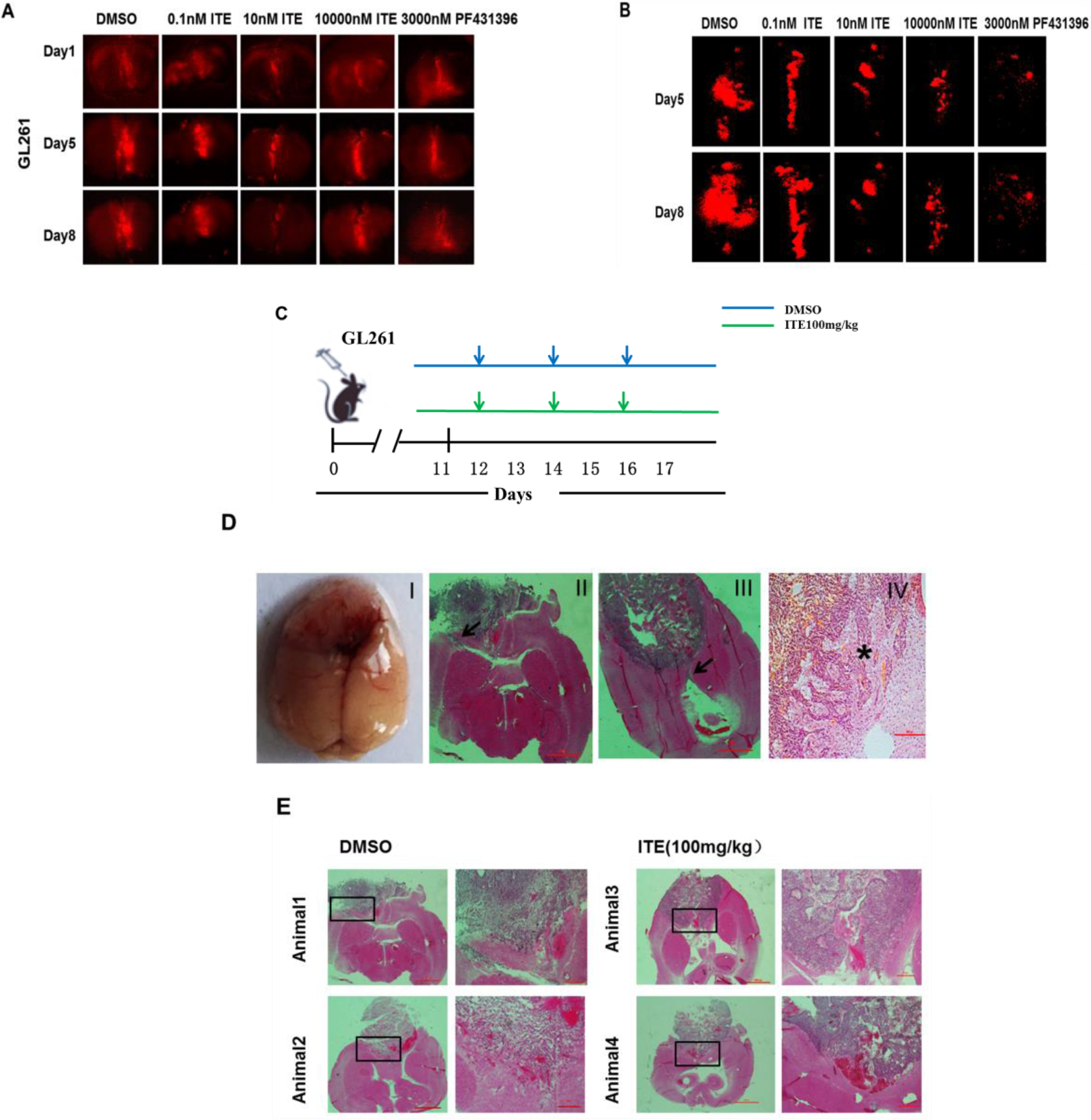
ITE blocked glioma cell invasion into the brain parenchyma in an ex vivo brain slice model and in an orthotopic mouse model of glioma. (A) Whole brain images of labeled GL261 cells invade into cultured mouse brain slices. (B)Cells were incubated with CM-Dil Dye, seeded into the seam of two adjacent brain slice halves cultured at the air-liquid surface with DMSO, a various dosage of ITE, or FAK inhibitor PF431396. Representative whole brain slice images were obtained on day1, day5, and day8. High-resolution brain slice culture images obtained by stitching overlapping images. (C) Experimental design and treatment schema. (D) Glioma invasion observed in the orthotopic mouse model (I) Gross appearance of a mouse brain with a glioma in the cerebrum. (II-III) Representative images of whole-brain horizontal frozen sections with H&E staining showing glioma invasion toward the ventricles. (IV) Representative image showing glioma invasion into the brain parenchyma. Arrows, ventricles. Asterisk, invading tumor cells at the tumor edge. (E) Invasion patterns of mice treated with ITE compared to DMSO. Tumor-bearing mice were treated with DMSO or 100mg/kg body weight ITE, and were euthanized. Horizontal brain sections from two animals per group were shown.

### ITE inhibited mesenchymal migration of glioma cells in 3D collagen matrix

By plating the cells in the type l collagen matrix, we observed various morphology (Video. 3A-F). One-hour ITE treatment led to the more spreading cell body and more filopodia extension and retraction (Video. 3A-B). In the control group, when a prominent lamellipodia-like protrusion detached, a shake of the extracellular matrix was observed (Video. 3A), indicating a strong attachment, which was not found in the ITE treated group (Video. 3B). Furthermore, the prevalent protrusion of ITE treated cells have more vibrant ruffles at the tip, confirming an unstable attachment to the ECM. Efficient mesenchymal migration was observed in the control group cells, overcoming constrictions in the matrix (Video. 3C).However, in the ITE treated cells, the cell rear remains attached to the ECM while its front part moves forward, resulted in cells with an elongated rear (Video. 3D), indicating compromised cellular contractility.

### ITE inhibited amoeboid migration of glioma cells

In the collagen matrix, we observed mesenchymal migrating cells started to bleb when encounter matrix constrictions, indicating that cells could swiftly switch to amoeboid migration (Video. 3C). Moreover, the A1 mode of the amoeboid migration, characterized by a prominent leading protrusion and round cell shape was also observed[9](Video. 3E). No typical amoeboid migration was observed in the ITE treated cells (Video. 3F), only a few spherical non-motile cells that continuously bleb, lacking sign of cellular polarization such as a stable large bleb on one side or multiple dynamic blebs on all but one side of the cells. Accordingly, ITE treated cells in 2D culture also failed to undergo amoeboid migration when subject to pH changes. When taken out of the incubator for 2hr photography, the culture medium became basic and U87MG cells responded to the pH change by becoming spherical and migrating, which is evident in the overlapping photos taken at the beginning and end of the photography period (Fig. 3G, Video. 3I). ITE treated cells remained in the same position and maintained the same shape, failed to respond (Fig. 3H, Video. 3J). There are a few spherical cells in the ITE group, but they also lacked the sign of cell polarity.

**Fig. 3.**
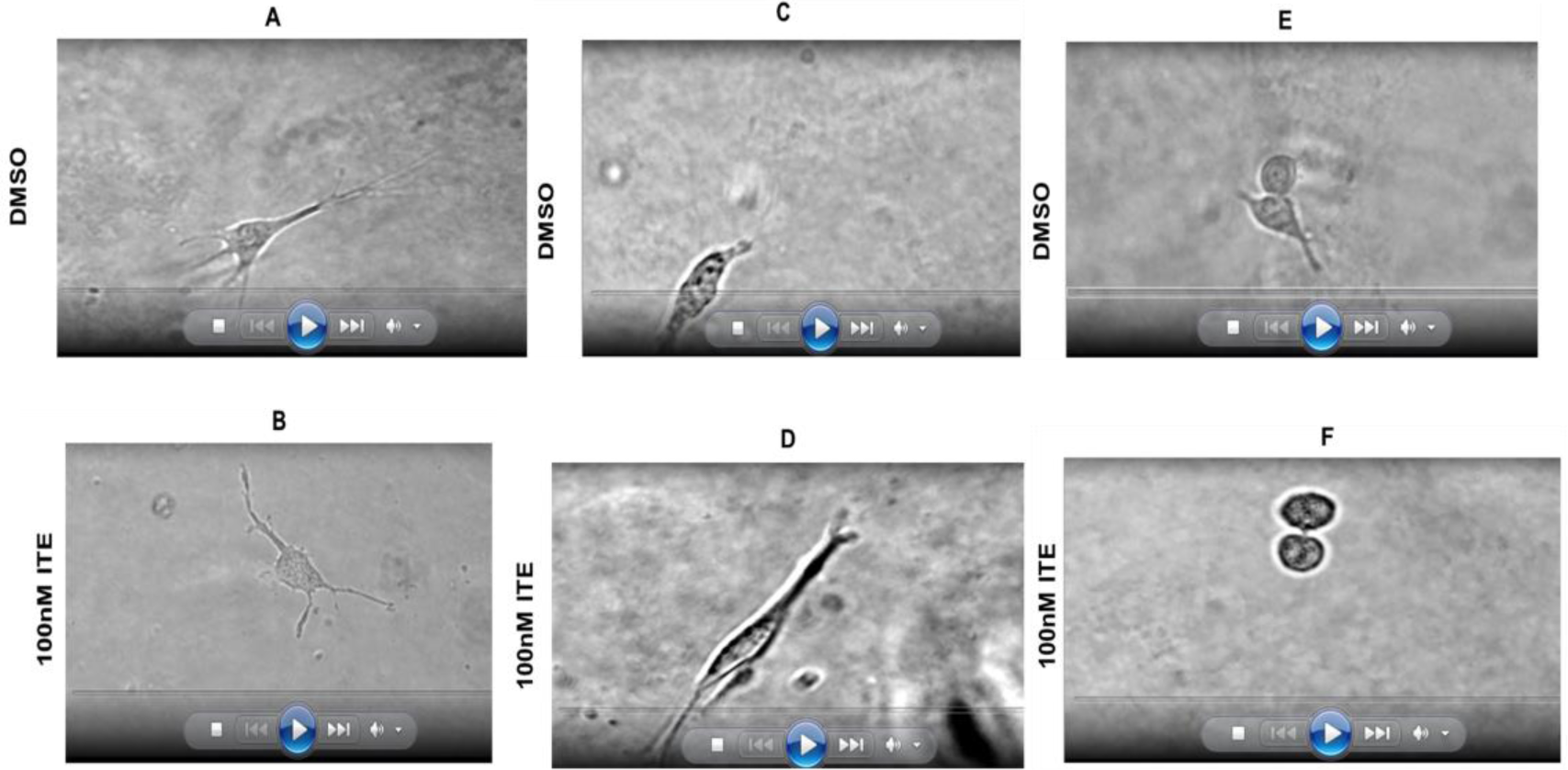
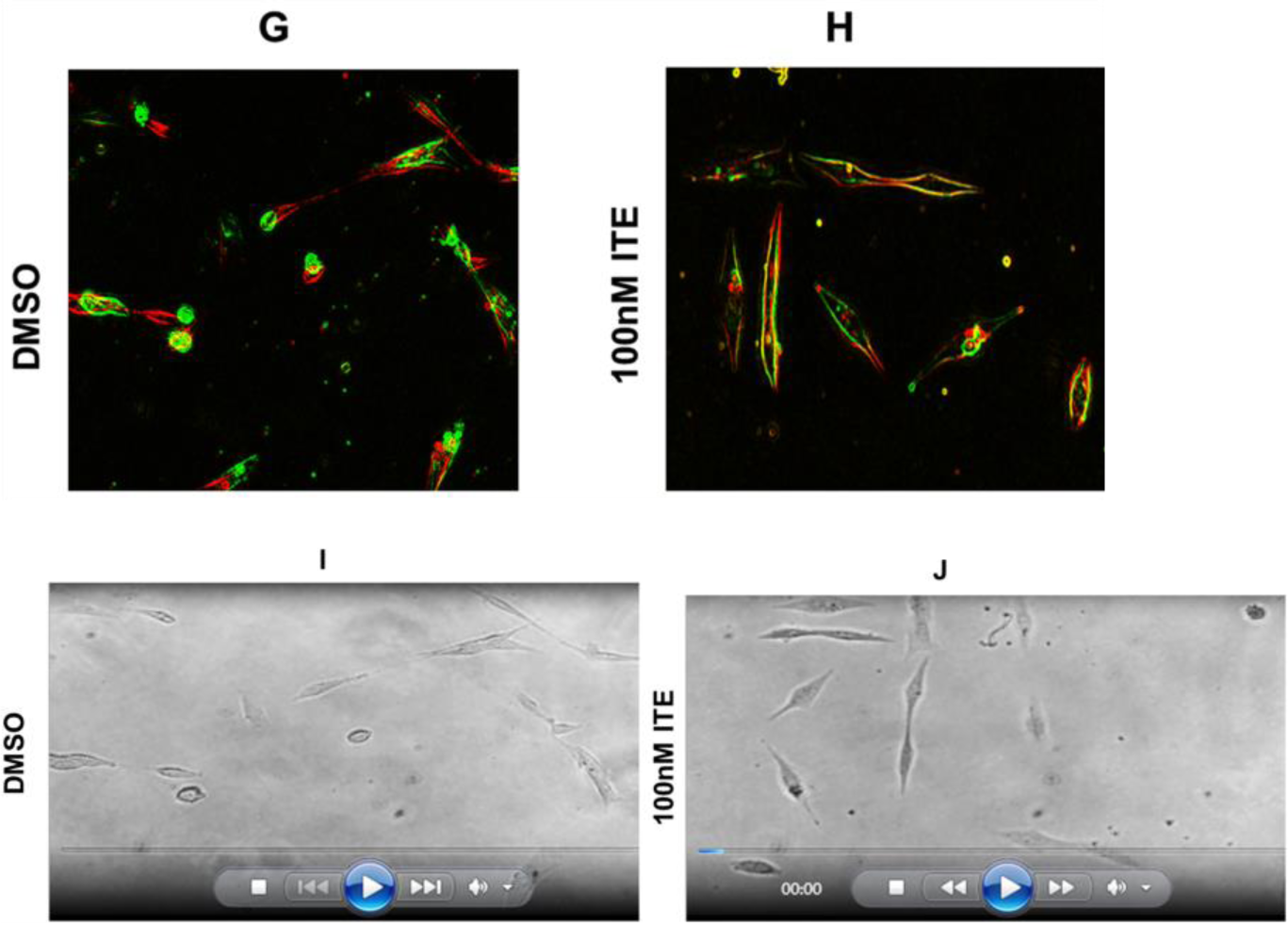
Effects of ITE on the spreading, protrusion attachment, and migration of glioma cells in 2D and 3D collagen matrix. (A-F) U87MG cells were plated in the type I collagen matrix, settled overnight, treated with either ITE or DMSO, and recorded by time-lapse photography: (A-B) Cells were treated with DMSO or ITE for 1hr. (C-D) Cells were treated with DMSO or ITE for 2hr. (E-F) Cells were treated with DMSO or ITE for 3hr. (G-J) U87MG cells in 2D culture were treated with DMSO or ITE for 18hr, recorded for 2 hr. (G-H) Overlap of photos taken at the beginning (Red Fluorescence) and end of the 2 hr (Green Fluorescence). (I-J) Videos of U87MG cells in 2D culture.

### MYH9 mediated ITE migration-inhibiting effects in human glioma cell

Since ITE treated cells display cytoskeletal changes, we studied related molecules. Several integrins, collagens, focal adhesion kinase PTK2 were examined by either RT-PCR or western blot, and none showed significant changes (Supplemental Figures). MYH9, a component of the non-muscle myosin IIA(NMIIA), was found to be significantly reduced by ITE treatment, both by RT-PCR and western blot analysis (Fig. 4A-B). To confirm the relationship between ITE and MYH9, we over-expressed the MYH9 gene in the U87MG cell by transfection of MYH9 cDNA fused to mCherry. Imaging and RT-PCR showed the elevated MYH9 level in transfected cells (Fig. 4C-D). Over-expression of MYH9 abrogated 0.1nM and 1nM ITE’s migration-inhibiting effects in human glioma cells (Fig. 4E-F), indicating that ITE’s migration-inhibiting effects were mediated by MYH9 at least partly. A strong correlation between MYH9 mRNA level and the area that cell migrated was observed in both the control group and over-expressing MYH9 (Fig. 4G), which reinforced the relationship between MYH9 expression and migration ability in the ITE treated cells.

**Fig. 4.**
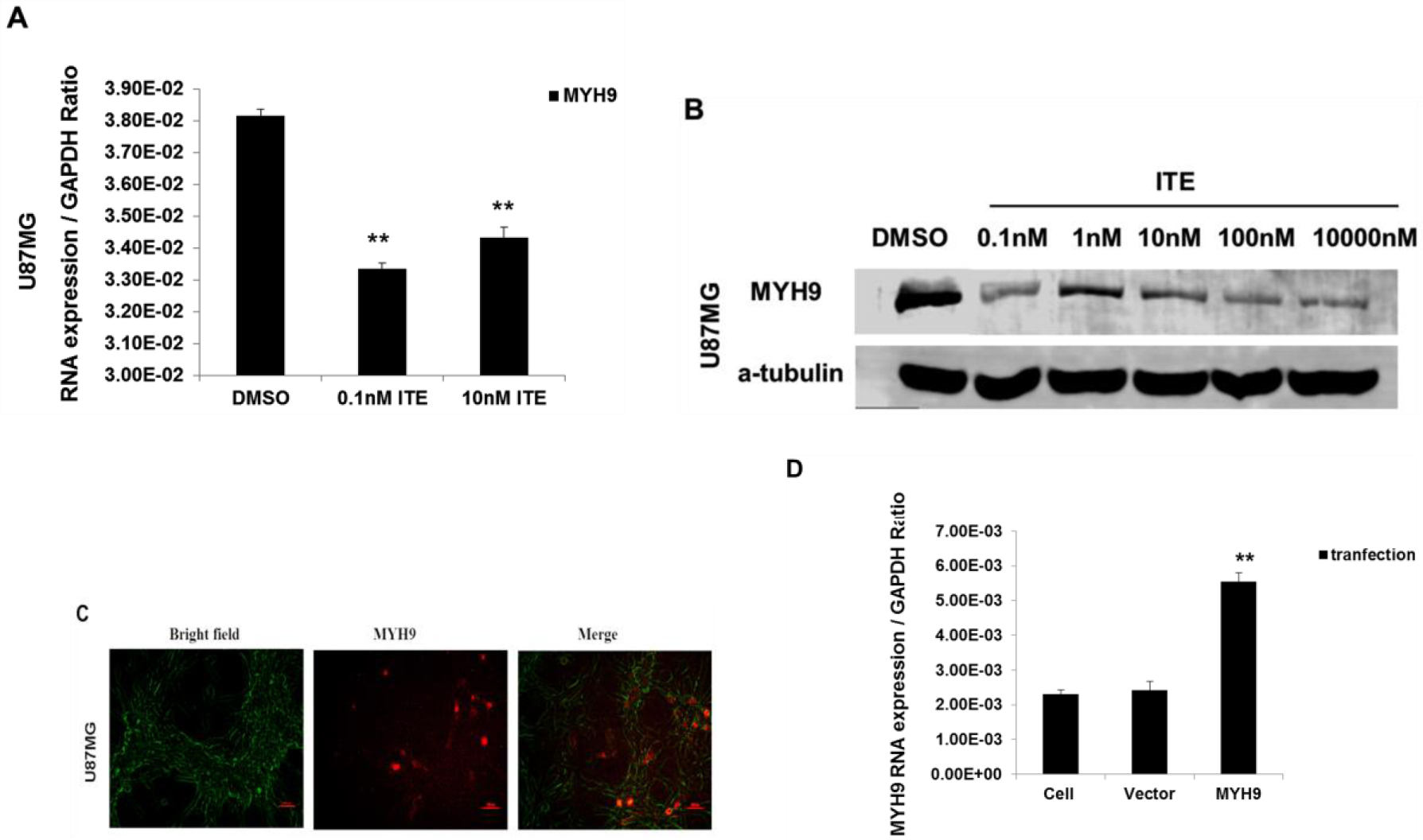
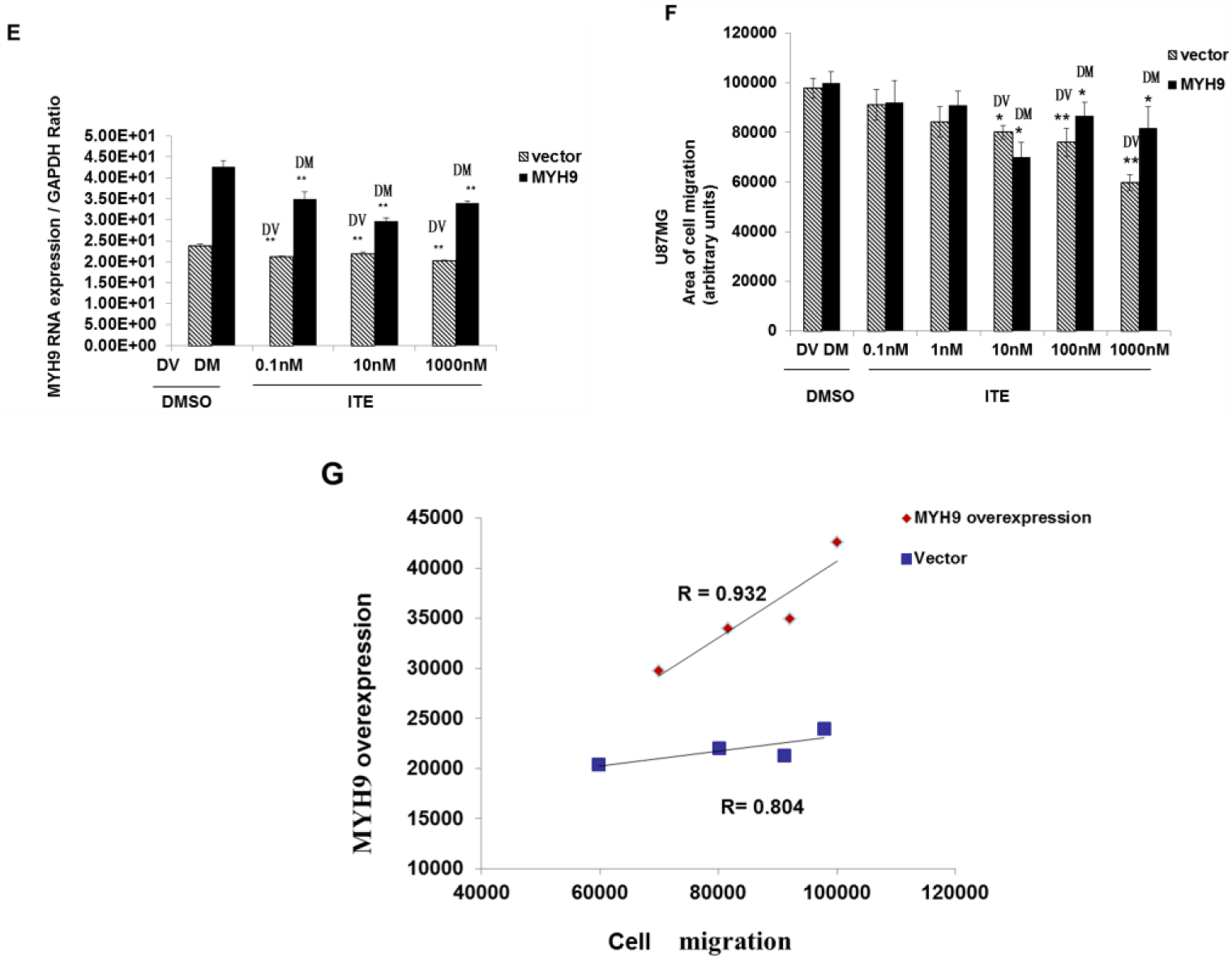
MYH9 expression level and cell migration ability. (A) Cells were treated with ITE for18hr and mRNA levels of MYH9 were determined by RT-PCR using GAPDH as a reference gene. (B) MYH9 protein levels were assessed in U87MG by western-blot. Cells were treated with various doses of ITE or DMSO. The bar graph summarizes the semi-quantitation of the western blot data. (C-D) Over-expression of MYH9-mCherry fusion protein by transfection of the U87MG cells. Red fluorescence was observed in cells 36hr after transfection with the MYH9 plasmid. MYH9 mRNA level was determined by RT-PCR using GAPDH as a reference gene (**p<0.01 vs control group) in MYH9 transfected cells, with an empty vector or non-transfection cells as control. (E) ITE reduced MYH9 mRNA level in U87MG cells over-expressing MYH9. MYH9 plasmid or empty vector-transfected cells were treated with either ITE or DMSO for 18hr and MYH9 mRNA levels were determined. (F) ITE’s effect on the migration ability of the U87MG cells that over-expressing MYH9. Either MYH9 plasmid or empty vector-transfected cells were treated with various doses of ITE or DMSO for 20hr. The area of cell migration was measured with 5 replicate wells per treatment condition (*p<0.05**P <0.01). (G) Correlation between MYH9 mRNA level and cell migration area in the U87MG cells.

## Discussion

Drugs that can block cancer cell invasion are urgently sought after, and we discovered that ITE, an endogenous ligand of AHR, could block migration of the glioma cells *ex vivo* and *in vivo*, supporting ITE to be investigated as a lead compound. We identified a previously unknown target of AHR, MYH9, providing a mechanism of how AHR regulates migration in human cells. MYH9 is essential for both mesenchymal and amoeboid migration, and accordingly, we observed inhibition of both migration modes in ITE treated glioma cells *in vitro*.

In comparison to other AHR ligands, ITE affected a different target with a higher efficacy on blocking cell migration. AHR ligands that have been reported for inhibiting cell migration include quercetin, omeprazole, indirubin, 3-methylcholanthrene(3-MC), etc. These ligands’ effective concentration varies from several to hundreds of micromolar [10–13]. ITE inhibits glioma cell migration at nanomolar concentrations, exhibiting a significantly higher efficacy. Various migration-related pathways were regulated by ligand-bound AHR, including phospholipase D1, CXCR4, PTK2, etc. We found that ITE inhibited migration and invasion via the AHR-MYH9 axis, supporting MYH9 as an AHR target.

Most of the observed U87MG cell behavior upon ITE treatment could be explained by weakened actomyosin contractility due to lowered MYH9/NMIIA level. Cell spreading and protrusion dynamics are the net results of the centrifugal force generated by actin polymerization and the centripetal force generated by actomyosin contractility[14, 15], so with lower actinomyosin contractility resulted from lower MYH9, both cells spreading and filopodia extension increased, especially at the lateral side of the cell, as NMIIA inhibited lateral filopodia formation and maintained cell polarity. The weakened protrusion tip attachment probably resulted from lowered NMIIA too, as inhibition/knockdown NMIIA resulted in reduction of the filopodia attachment strength[16], and inefficient lamellipodia attachment caused the ruffles [17]. Cell rear retraction depends on myosin IIA, which explained the inefficient rear retraction upon ITE treatment[18]. All these changes are similar to what was reported in cancer cells with depleted or lowered NMIIA[19, 20].

The inhibition of the mesenchymal to amoeboid transition in both 2D and 3D culture could have resulted from the lower MYH9 expression level upon ITE treatment. Cells need actomyosin contractility to become spherical[21], hence most of the ITE treated cells remained in its original polygon shape. The inefficient blebbing amoeboid mode manifested as a lack of polarity is probably also due to the lower MYH9, which is needed to form cell rear[22]. Blebs forming and retracting continuously in the round amoeboid migration, yet there is a lack of blebbing at the rear, which is essential for cell polarity.

Among the four possible cancer cell migration modes, we discovered that ITE inhibited both mesenchymal and the characteristic amoeboid A1 migration mode[23]. In GBM, to migrate in the brain, glioma cells have to squeeze through pores smaller than its cell body size, often with the help of matrix metalloproteinase where NMIIA is essential [24, 25], too. The reason for the lack of A1 mode in ITE treated group needs further investigation. Moreover, MYH9 promotes tumor progression by modulating the immune micro-environment, so reduced MYH9 level by ITE treatment potentially offers immune-promoting benefit in addition to its invasion-inhibiting effects [26]. When writing this paper, it is reported that Programmed death-ligand1 (PDL1)/Programmed cell death protein 1(PD1) blockade was effective as neoadjuvant therapy in GBM. The combination of ITE and PD1/PDL1 blockade warrants further research.

## Conclusion

Small molecular AHR agonist ITE regulated glioma cells’ migration via binding to AHR and inhibited the actinomyosion contraction due to reduced MYH9 protein level. ITE efficiently blocked glioma invasion in vitro, ex vivo, and in vivo in an orthotopic mouse glioma model.

## Supporting information

Several integrins, collagens, focal adhesion kinase PTK2 were examined by either RT-PCR or western blot

## Acknowledgments

This work was supported by the National Natural science Foundation of China [grant 81460455]. We thank Dr. Jun Dong [Suzhou University]kindly taught us how to build the orthotopic glioma model and Professor Xiaomei Zhu from [Shanghai University] who provided insight and expertise that greatly assisted the research.

