## Supplementary material for "Targeting Aryl Hydrocarbon Receptor with small molecule, 1′H-indole-3′-carbonyl-thiazole-4-carboxylic acid methyl ester blocked human glioma cell invasion via MYH9": Several integrins, collagens, focal adhesion kinase PTK2 were examined by either RT-PCR or western blot

[**Supplementary Table 1**](manuscript.docx)**.** List of primary antibodies used for western blotting.

| **Antibodies** | **Company** | **Catalog Number** | **Dilutions** |
| --- | --- | --- | --- |
| Mouse anti- AHR | Abcam | Ab-43199 | 1:1000 |
| Rabbit-anti-PTK2 | Cell Signaling | 9330 | 1:1000 |
| Rabbit-anti- MYH9 | Proteintech | 14844-1-AP | 1:500 |
| GAPDH | Proteintech | 10494 | 1:2000 |
| Mouse-anti-alpha tubulin | Proteintech | 66031 | 1:2000 |
| IRDye 800CW Goat anti-Mouse secondary antibody | LI-COR | 925-32210 | 1:15000 |
| IRDye 800CW Goat anti-Rabbit secondary antibody | LI-COR | 925-32211 | 1:15000 |

[**Supplementary Table 2**](manuscript.docx)**.** List of sequences for primers of genes.

| Gene (human) | Forward primer | Reverse primer |
| --- | --- | --- |
| PTK2(human) | ACATTATTGGCCACTGTGGATGAG | GGCCAGTTTCATCTTGTTGATGAG |
| HPRT(human) | TGACACTGGCAAAACAATGCA | GGTCCTTTTCACCAGCAAGCT |
| MYH9(human ) | ATCTCGTGCTATCCGCCAAG | GTTGTACGGCTCCAACAGGA |
| GAPDH(human) | TCATTGAGCCCTTCCACAATG | GGTGTGAACCACGAGAAATATGAC |
| COCL1(human) | TCTGCGACAACGGCAAGGTG | GACGCCGGTGGTTTCTTGGT |
| ITGA5(human) | GCCTGTGGAGTACAAGTCCTT | AATTCGGGTGAAGTTATCTGTGG |
| ITGB5(human) | CAGGTGGAGGACTATCCTGTG | GTGCCGTGTAGGAGAAAGGAG |

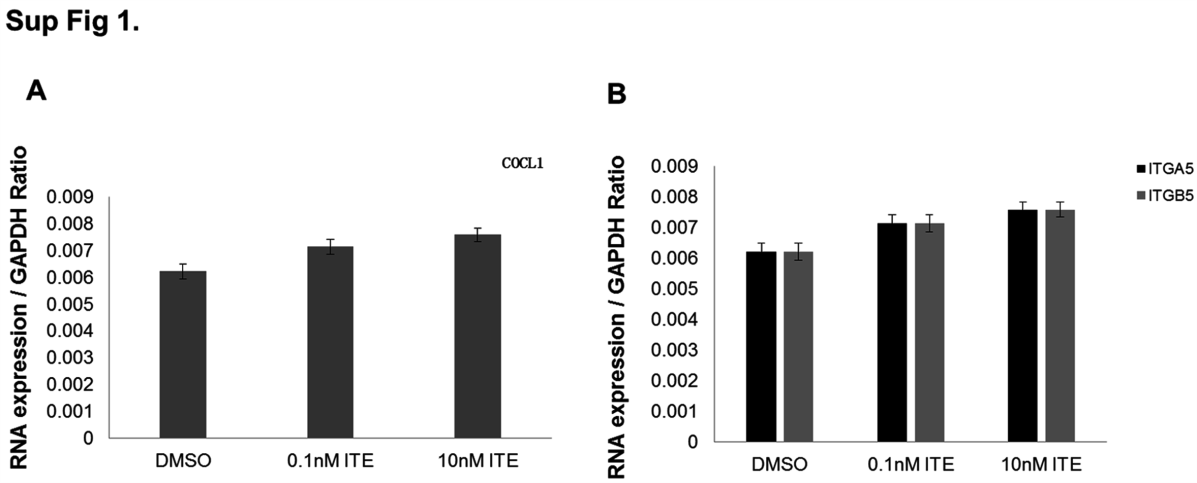

**Sup Fig1.**

ITE treatment detects integrin, collagen and other related genes’ expression level. A and B: cells were treated with ITE (0.1nM, 10nM) or DMSO (control) for 18h and mRNA level of COCL1 and intergrin (ITGA5,ITGB5) using HPRT as the reference gene.

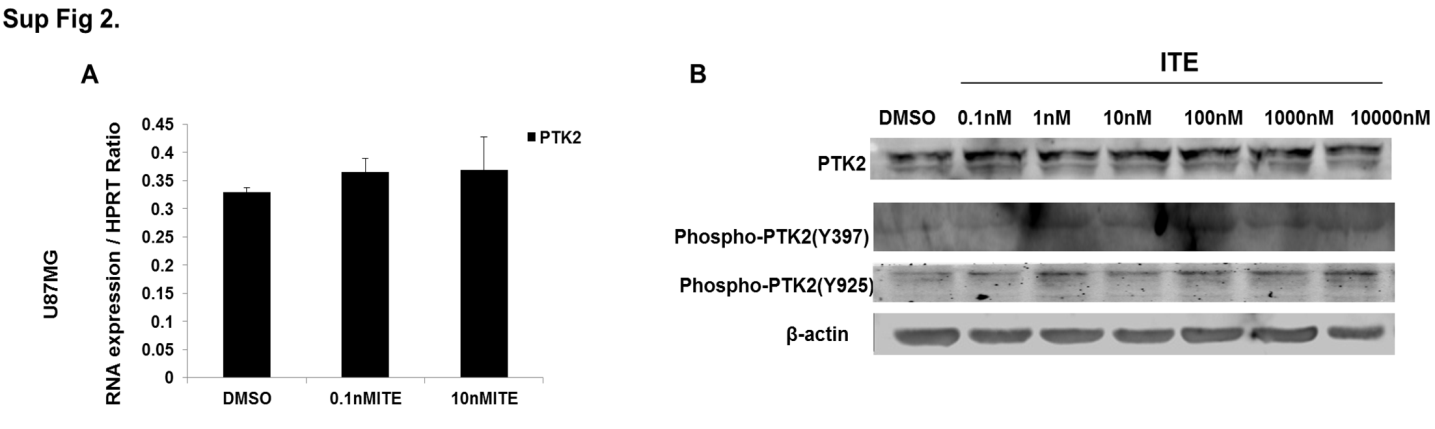

**Sup Fig.2**

ITE treatment detects PTK2 levels human glioma cells.（A）cells were treated with ITE (0.1nM, 10nM) or DMSO (control) for 18h and mRNA level of PTK2 using HPRT as the reference gene. B: PTK and phospho PTK2 protein levels were observed in treated U87MG by western-blot, cells were treated with ITE (0.1nM, 1nM, 10nM, 100nM, 1000nM,10000nM) compared with DMSO, and detect the PTK2 protein level.
